# Anticancer potential of piericidin A1 and derivatives isolated from *Streptomyces* sp. associated with *Palythoa variabilis* from Brazilian Reefs

**DOI:** 10.1101/2025.11.19.689357

**Authors:** Bianca Del B. Sahm, Katharine G. D. Florêncio, Francisco C. L. Pinto, Ana I. V. Maia, Carlos A. M. Rocha, Paula C. Jimenez, Otília D. L. Pessoa, Tito M. C. Lotufo, Leticia V. Costa-Lotufo, Diego V. Wilke

## Abstract

Marine holobionts are prolific sources of bioactive natural products. Previously, our group investigated cytotoxic compounds in zoantharians of the genus *Palythoa* and, herein, we further explored the anticancer potential of marine bacteria associated with *P. variabilis*. After isolating 10 culturable bacterial strains, we screened for cytotoxic activity on metastatic prostate tumor cells (PC-3/M) using the MTT assay. The crude extract of one of the strains (BRA-035), identified as *Streptomyces* sp., was selected due to its bioactivity profile and submitted to cytotoxicity-guided fractionation. This approach allowed us to isolate piericidin A1 and identify other 2 piericidin derivatives. Piericidin A1 displayed a wide range of potency, with IC_50_ varying from pM to low nM for several tumor cells (OVCAR, PC-3, PC-3/M, HCT-116) or non active (IC_50_ >12uM) for leukemia cell line (HL-60) and murine melanoma (B16-F10). Overall, our findings demonstrate that *P. variabilis* hosts bacteria with biotechnological potential, confirming piericidin A1 as a promising candidate for anticancer applications.

## Introduction

Zoantharians (Cnidaria: Anthozoa: Hexacorallia) are soft-bodied benthic marine animals commonly known as sea mats that form colonies covering rocky shores and various reef habitats worldwide. They comprise a taxonomically diverse group, which includes the genera *Palythoa, Zoanthus*, and *Parazoanthus* ^[1,2]^. Zoantharians present specialized metabolic pathways that allow them to survive in hostile environments, providing competitive advantages in space occupation, food acquisition, and defense against predators. Beyond their species diversity, they are also recognized for their rich chemical arsenal with notable biological activities, which continues to motivate natural product research and the development of novel pharmaceutical agents ^[3]^.

Like other cnidarians, zoantharians harbor an intrinsic microbiota that plays crucial roles in maintaining homeostasis and overall health, framing these animals as holobionts ^[4,5]^. In addition to these essential functions, bacterial communities associated with zoanthids are now recognized as key contributors to the production of specialized metabolites. Investigations into the biosynthetic machinery of invertebrates have revealed microbial participation in the synthesis or modification of bioactive compounds. This has led to the hypothesis that many substances previously attributed to marine invertebrates are, in fact, produced by their symbiotic or associated microorganisms ^[6]^. Indeed, studies exploring the biosynthetic potential of marine bacteria — both associated with holobionts and free-living, using cultivation-dependent and-independent approaches — have uncovered new chemical classes, derivatives, or analogues with significant biological activity against pathogens and diseases, including cancer ^[7–10]^.

Most drugs originated from natural products ^[11]^. From a sustainable development perspective, drug discovery programs focused on natural products are strategic due to their legacy of yielding for deriving new medicines and, furthermore, to support biodiversity conservation and equitable profit sharing. Therefore, besides being a promising source for the development of new therapeutic entities, this approach is also connected to a highly demanded appeal for conservation of ecosystems in a climate change scenario ^[12]^. In this context, marine natural products (MNP) serve as an important source of bioactive molecules, despite their recent history compared to terrestrial counterparts ^[13]^. Underlining the pharmacological potential of MNP, 20 drugs are currently in clinical use, mainly as antineoplastics ^[14]^.

Our group has systematically investigated bioactive molecules from marine organisms to highlight the therapeutic potential of Brazilian marine biodiversity ^[15]^. Previously, we have isolated structurally diverse compounds, including palyosulfonoceramides ^[16]^ and lipidic α-amino acids (LAAs) from *P. variabilis* ^[17]^. LAAs from *P. variabilis* displayed antiproliferative activity against tumor cell lines, inducing apoptotic features and DNA fragmentation ^[18]^. Later, two ergostane-type sterols were isolated from *P. variabilis* and *P. caribaeorum*, and also showed growth inhibition against colorectal cancer cells ^[19]^. Chromomycin A (CA_5_), along with 3 new chromomycins – CA_6_ CA_7_ and CA_8_ – isolated from the extracts of *Streptomyces* sp. BRA-384 associated with *P. caribaeorum* was also identified ^[20]^ and further studied on its anticancer potential. Chromomycins are glycosylated aureolic acids known to bind to the DNA minor groove, causing double-strand DNA damage and triggering cell stress and death ^[21,22]^. Our studies revealed that CA_5_ binds to the transcription factor Tbx2 and inhibits melanoma cell proliferation^[22,23]^. In the present study, we screened the cytotoxicity of crude extracts obtained from bacteria associated with *P. variabilis* from Northeastern Brazil and identified the active compounds. We also discuss the pharmacological properties of piericidin A, a highly cytotoxic compound with anticancer potential isolated from bacteria associated with the zoantharian.

## Results and Discussion

### Isolation of bacteria associated with *Palythoa variabilis* and bioactivity screening

Initially, 10 bacteria strains associated with *P. variabilis* collected at Taiba beach rocks were isolated and fermented in A1 broth for 8 days under agitation (SI.1). The activity of crude ethyl acetate extracts obtained from isolated bacteria was tested on PC-3/M human prostate carcinoma cell line. As shown in Figure 1A, the crude extract obtained from BRA-035 strain caused 100% growth inhibition on metastatic prostate cancer cells (PC-3/M) at 50 µg/mL. The extracts obtained from other isolates did not show activity against these cells. BRA-035 colonies depict a dry, brownish appearance, and produce white spores within 10 days on A1 agar medium at 28°C (Figure 1B). Molecular identification by 16S rDNA sequencing analysis suggests the BRA-035 strain belongs to the *Streptomyces* genus (family *Streptomycetacea*; order *Actinomycetales*), and will be referred to, from now on, as *Streptomyces* sp. BRA-035 (Figure 1C). The *Streptomyces* genus is known to be an excellent producer of metabolites with biological activity. Indeed, these bacteria are recognized as producers of a great number of clinically relevant pharmacological substances with a wide range of activities, such as antimicrobial (tetracycline, neomycin, streptomycin), antifungal (amphotericin B, nystatin, natamycin), anticancer (doxorubicin, daunorubicin, mitomycin C) and anthelmintic (avermectin) ^[24–27]^.

**Figure 1.**
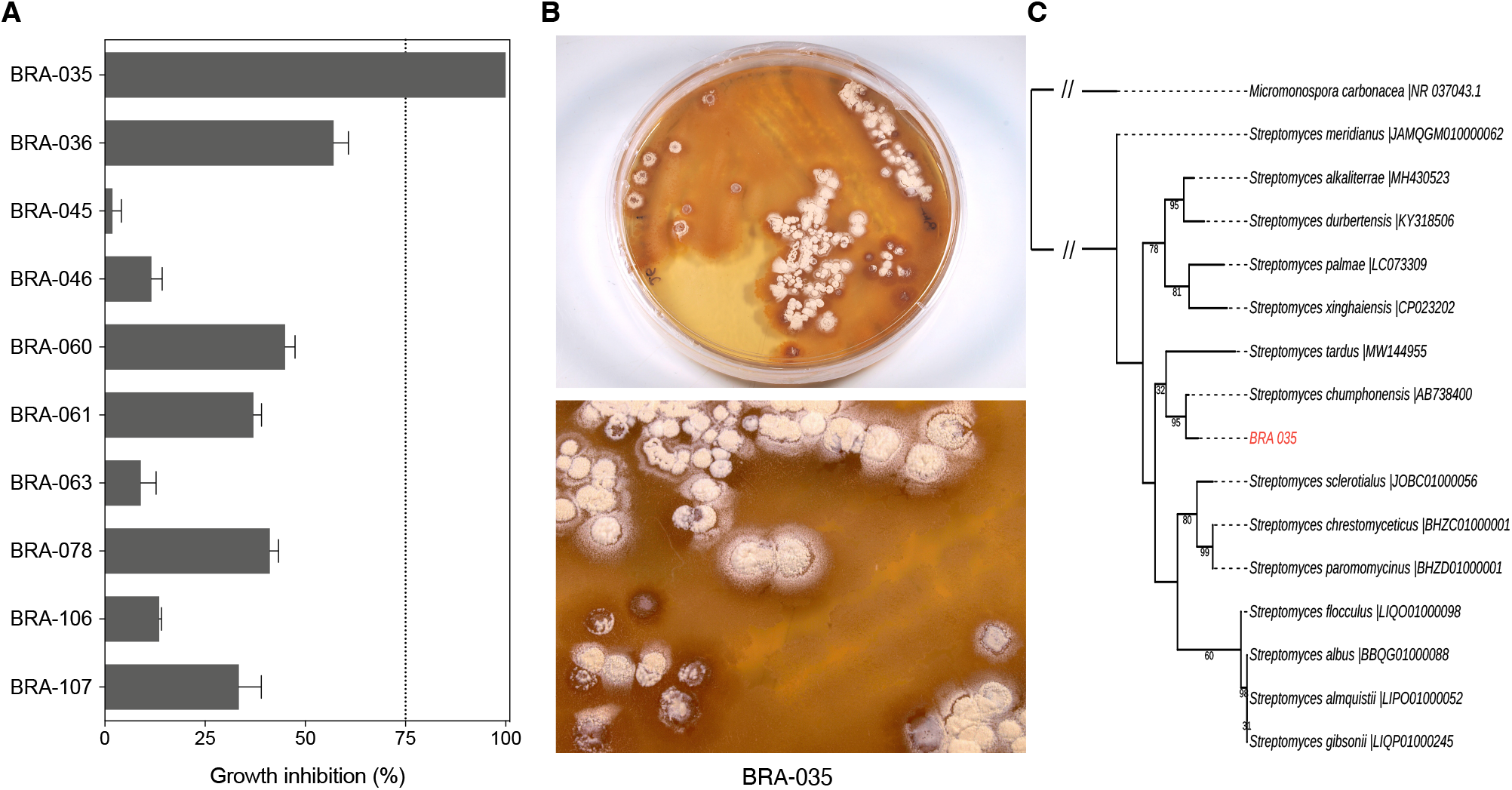
The BRA-035 strain isolated from the zoantharian *Palythoa variabilis* exhibited the highest inhibition of tumor cell growth. **A**, Bar graph showing the percentage (mean+SEM) of growth inhibition of metastatic prostate cancer cells (PC-3/M) after 72 h of incubation with crude extracts from bacterial isolates associated with *P. variabilis*, determined by the MTT assay. **B**, Images of the BRA-035 strain grown on A1 agar media. **C**, Phylogenetic tree based on 16S rRNA gene sequences showing the relationship between BRA-035 and closely related *Streptomyces* species.

### Bioassay-guided isolation of bioactive compounds of *Streptomyces* sp. BRA-035

In order to identify the bioactive substances produced by the *Streptomyces* sp. BRA-035, 10L of A1 culture broth was cultivated for 8 days, yielding 150 mg of crude extract. The extract was fractionated using high performance liquid chromatography (HPLC). The cytotoxicity of the 8 fractions obtained (SI.2) was evaluated on PC-3/M cells by MTT assay for 72h. As shown in Table 1, the crude extract and fractions F7 (BRA-035-F7) and F8 (BRA-035-F8) showed potent cytotoxic activity with inhibition concentration mean (IC_50_) of 0.48 ng/mL, 47 ng/mL, and <16 ng/mL, respectively, against PC-3/M cells, while the remaining fractions were considered inactive (SI.3 and SI.4). Further analysis of fractions F7 and F8 led to the characterization of a purified compound (SI.5 and SI.6).

**Table 1.** Cytotoxicity of BRA-035-*Streptomyces* crude extract and fractions on metastatic prostate cancer cells. Results shown as inhibition concentration mean (IC_50_), and confidence interval of 95% (CI95%) on PC-3/M cell line after 72h incubation. Data from three experiments performed in triplicate were determined by non-linear regression.

| Sample | IC<sub>50</sub> (ng/mL) | CI95% |
| --- | --- | --- |
| BRA-035 crude extract | 0.046 | –<sup>a</sup> |
| BRA-035-F1 | > 50000 | –<sup>a</sup> |
| BRA-035-F2 | > 50000 | –<sup>a</sup> |
| BRA-035-F3 | > 50000 | –<sup>a</sup> |
| BRA-035-F4 | > 50000 | –<sup>a</sup> |
| BRA-035-F5 | > 50000 | –<sup>a</sup> |
| BRA-035-F6 | > 50000 | –<sup>a</sup> |
| BRA-035-F7 | 46.90 | 23.19 - 94.55 |
| BRA-035-P8 | < 0.016 | –<sup>a</sup> |

<sup>a</sup> - Not determined by non-linear regression

The molecular formula C_25_H_37_NO_4_ (nine degrees of unsaturation) of BRA-035-F8 was determined by the high-resolution mass spectrometry (HRESIMS) from the protonated molecular ion observed at *m/z* 416.2848 [M+H]^+^ and the ion at *m/z* 398.2560, which indicated the loss of a water molecule [M+H-H_2_O]^+^ (SI.7). Its ^1^H NMR spectrum (SI.8) showed signals to olefinic protons at *δ*_H_ 6.10-5.20, two methylenes *δ*_H_ 3.41 and 3.80; two methoxyl groups *δ*_H_ 3.87 and 3.99, two methines *δ*_H_ 3.62 and 2.70, and six methyl groups at *δ*_H_ 2.10-0.80. Comparison of the data with those of piericidin A1 (**1**) showed a good match ^[28–31]^. The chromatogram of BRA-035-F7 (SI.5B) shows in the LC-MS, a peak with retention time (t_R_) at 24.50 min, corresponding to the molecule protonated [M+H]^+^ a ion at *m/z* 578.3431, indicating the molecular formula C_31_H_47_NO_9_ (ten degrees of unsaturation). The mass fragment at *m/z* 398.2710 (SI.9) indicated the loss of the glucose moiety [M+H-Glu]^+^ leading us to suggest the structure of **2** as the glucopiericidin A1 (**2**) ^[32,33]^. Another compound with t_R_ 26.8 min (SI.10) and *m/z* 432.2808 [M+H]^+^ and [M+H-MeOH]^+^ suggested the molecular formula of C_25_H_37_NO_5_. Based on the mass fragmentation, the compound **3** was identified as piericidin C1 (**3**) ^[33,34]^.

Piericidins A and B were first isolated from *S. mobaraensis* in 1963 for insecticidal application ^[30]^. Later, the antimicrobial and antitumour potential of piericidins, including **1** and **2**, were reported ^[32,35–37]^. Most known piericidins were obtained from marine and terrestrial bacteria of the genus *Streptomyces*, with a few exceptions occurring in other genera within Actinomycetota phylum ^[38]^. To corroborate this information, here we describe the isolation and identification of three piericidins produced by a *Streptomyces* strain recovered from *P. variabilis*. The chemical structure of **1** consists of a 2,3-dimethoxy-5-methyl-4-pyridinol ring coupled at C-6 with a poly-methylated and - insaturated side chain ^[39]^ (Figure 2). The biosynthetic pathway of piericidin A1 was unraveled from the sequencing of the complete genome of *S. piomogeues* ^[31]^, which showed six modular polyketide synthases, two methyltransferases, two amidotransferases - one of which is an ATP-dependent aminotransferase for the N atom, and a monooxygenase are required to produce the substance. Some authors suggest that other natural piericidin aglycones (i.e. piericidin C1) have apparent production during the fermentative growth process or extraction phases ^[28]^, while glycosidic derivatives are produced by glycosylation of piericidin A1 as a consequence of the long period of growth in culture medium ^[32,38]^.

**Figure 2.**
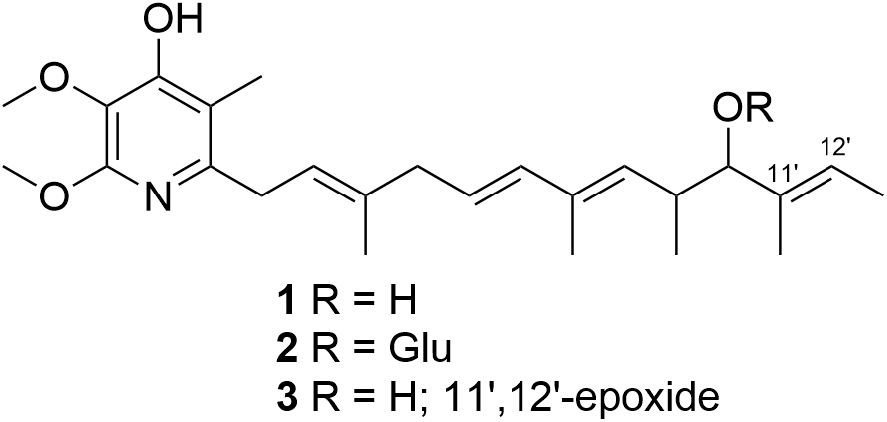
Chemical structure of piericidins isolated from *Streptomyces* sp. BRA-035: Piericidin A (**1**), glucopiericidin A1 (**2**) and piericidin C1 (**3**).

### Effect of piericidin A1 on viability of cancer cells *in vitro*

The effects of **1** on cell viability of seven tumor cell lines were investigated by MTT assay. Compound **1** showed unprecedented high potency against OVCAR-8, PC-3/M, HCT-116, SF-295 and PC-3 cell lines, with IC_50_ values ranging from 500 fM (OVCR-8) to 9.0 nM (PC-3) (Table 2 and SI.11). In turn, HL-60 and B16-F10 were found to be resistant to **1** (IC_50_ >12uM). These unprecedented IC_50_ values for piericidin A1 obtained in this study highlight the importance of screening programs, including further investigation of compounds previously studied. Piericidins present a 4-hydroxypiridine nucleus (cyclic head) followed by a branched unsaturated side chain (hydrophobic tail), which is structurally correlated to ubiquinone Q_10_ (coenzyme Q), responsible for energy production through the mitochondrial respiratory chain ^[40,41]^. Indeed, piericidins are known to interfere with electron transport through the mitochondrial inner membrane in mammalian cells, specifically blocking the first step of the transport chain, known as the NADH-ubiquinone oxidoreductase (complex I) ^[28,29]^. Due to their structural similarity and physiological effect, piericidins can be classified as coenzyme Q antagonists, along with other complex I inhibitors, such as stigmatellin and rotenone ^[42]^. The consequences of blocking the mitochondrial respiratory chain with piericidins include reduced ATP production and increased ROS production. To a large extent, such energetic and reactive changes lead to cell death ^[43]^.

**Table 2.** Cytotoxic activity of **1** on a panel of tumor cell lines assessed by the MTT assay. Results are shown as mean inhibitory concentration (IC_50_), and 95% confidence intervals (CI95%) on tumor cell lines after 72h incubation. Data from three experiments performed in triplicate were determined by non-linear regression.

| Cell line | Tissue origin | Species | IC<sub>50</sub> (nM) | CI95% |
| --- | --- | --- | --- | --- |
| PC-3 | Prostate carcinoma | Human | 9.084 | 2.491 - 33.13 |
| PC-3/M | Metastatic prostate carcinoma | Human | 0.0018 | 0.0002 - 0.118 |
| HL-60 | Acute promyelocytic leukemia | Human | > 12000 | –<sup>a</sup> |
| HCT-116 | Colorectal carcinoma | Human | 0.144 | 0.062 - 0.338 |
| SF-295 | Glioblastoma | Human | 1.418 | 0.947 - 2.123 |
| OVCAR-8 | Ovarian adenocarcinoma | Human | 0.0005 | 0.0002 - 0.013 |
| B16-F10 | Metastatic melanoma | Murine | > 12000 | –<sup>a</sup> |

<sup>a</sup>– Not determined by non-linear regression

The process of tumorigenesis involves acquisition of key features to provide adaptive advantages for tumor cells’ proliferation, survival, and dissemination as much as possible ^[44]^. It is worth highlighting that tumor cell metabolism must enable uninterrupted cellular proliferation, which implies a high demand for energy to sustain these processes ^[45]^. This is particularly valid for cells cultured *in vitro*, which are kept in exponential growth to enable cytotoxic studies. The ATP generation via oxidative phosphorylation is the most efficient pathway; however, glycolysis also plays an important role in cancer cells ^[46]^. Glycolysis bypasses the oxidative phosphorylation pathway to overcome hypoxic conditions arising from fast proliferation rates. Known as the Warburg effect, glycolytic metabolism suppresses the oxidative pathway in some types of cancer cells, even in an aerobic environment ^[47–49]^. That being said, the contribution of mitochondrial respiration to energy production in malignant cells remains debated, and this pathway may represent a potential therapeutic target in cancer. In contrast to the widespread concept of glycolysis dependence, in cancer cells, the mitochondria oxidize several metabolic fuels by respiration to produce most of the cellular energy. Furthermore, several mitochondrial respiratory chain subunits are critical for cancer cell growth, metastasis, and invasion ^[50]^. The difference in sensitivity of the various cell lines assessed herein to **1** correlates with singularities in energy pathways. According to the literature, B16-F10 is highly dependent on glycolysis and produces high lactate amounts under aerobic conditions ^[51]^. HL-60 depicts high plasticity on energetic pathways, switching sources to adapt to different conditions ^[52]^. Indeed, these cell lines displayed remarkable resistance to **1** (Table 2). On the other hand, PC-3, PC-3/M, OVCAR, SF295, and HCT-116 are reported to have a greater dependence on oxidative phosphorylation to produce ATP ^[53,54]^, and showed greater susceptibility to **1**. This is a key point to explain the variation in sensitivity and, therefore, in the IC_50_ values obtained for **1** in the different cell lines, and deserves further investigation.

### Cell count and viability of piericidin A1 on tumor cells

It must be acknowledged that the blockage in the electron transport chain induced by compound **1** can impact the cell-viability measurements of piericidin A1-treated cells carried out by the MTT assay. The MTT assay is a widely used colorimetric method in drug screening programs. It provides a convenient and rapid approach to indirectly assess cell viability through their cellular metabolic activity ^[55]^. In this assay, MTT (3-(4,5-dimethylthiazol-2-yl)-2,5-diphenyltetrazolium bromide), a yellow tetrazolium salt, is reduced to insoluble purple crystals of formazan. Quantification is performed by measuring solubilized formazan spectrophotometrically ^[56]^. The reduction mechanism involves cleavage of the tetrazolium ring by dehydrogenase enzymes, and therefore depends on the cell’s metabolic activity driven by mitochondrial and cytosolic NADH flux ^[57]^. Therefore, it is clear that compound **1** interferes with cellular metabolism and may result in under-conversion of MTT to formazan, possibly caused by perturbations in oxidoreductase reactions ^[58]^.

To alternatively address the cytotoxicity profile of compound **1**, we resorted to a direct cell counting/cell viability method. Using flow cytometry to evaluate the effects of 1 on cell number and membrane integrity (cell death) after 24, 48, and 72h exposure, we selected one among the piericidin A1-sensitive (HCT-116) and resistant (B16-F10) cell lines from the panel initially assessed herein. HCT-116 cells incubated with **1** showed lower cell counts compared to the negative control (C-) at 24h, 48 and 72 h on concentrations ranging from 0.1aM to 12uM (Figure 3A). At 72h, cultures of HCT-116 cells exposed to 0.1 aM - 24 pM of **1** were reduced to nearly 50% of cell count of the control, while at 3.8 nM - 12 uM, that number came down to 25%. However, no change in cell viability was detected in most treatment conditions, except for a slight loss (∼20%) observed at 12 uM treatment after both 48 and 72h (Figure 3C). B16-F10, in turn, showed decreased cell count at 3.8 nM and 12 uM after 48h and 72h incubation, reaching nearly 50% and 40% cell counts after 72h exposure to **1**, respectively (Figure 3B). A slight decrease in membrane integrity of these cells was observed for the same concentrations and corresponding time periods (Figure 3D). These results indicate that, in fact, there is a marked difference in sensitivity to **1** between HCT-116 and B16-F10 cell lines, and, furthermore, that the MTT assay may underestimate the cytotoxicity of compounds that promote mitochondrial uncoupling.

**Figure 3.**
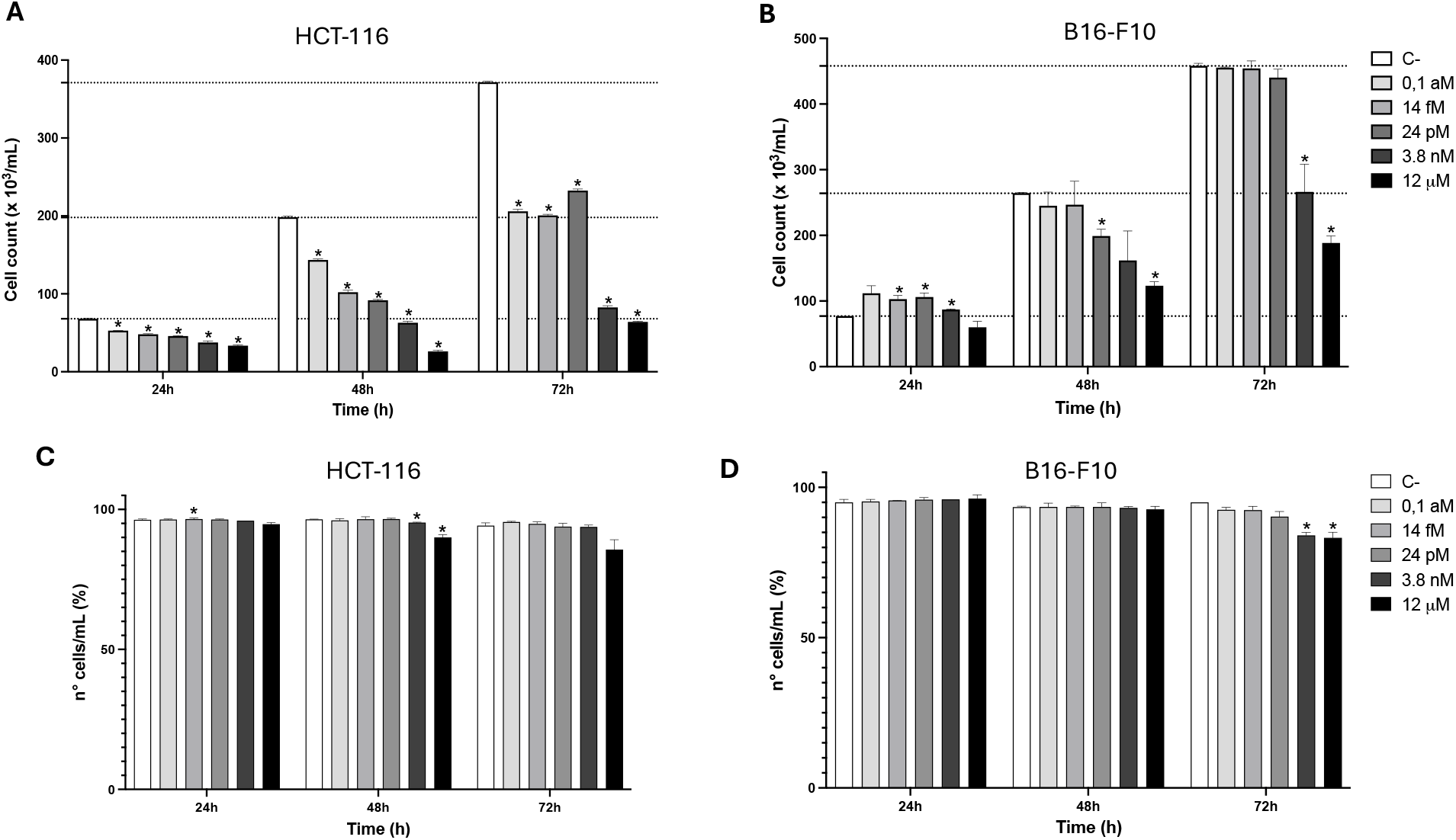
Effect of compound **1** on cell count and viability of HCT-116 and B16-F10 cell lines under different conditions. **A**, cell count of HCT-116 after 24, 48 and 72 h of treatments. **B**, cell count of B16-F10 after 24, 48 and 72 h of treatments. **C**, cell viability of HCT-116 after 24, 48 and 72 h of treatments. **D**, cell viability of B16-F10 after 24, 48 and 72 h of treatments. Data represent the mean+SEM of three independent experiments performed in triplicate. **\*** *p < 0*.*05*.

As observed for compound **1**, the role of metformin on inhibition of tumor growth is also related to the impairment of mitochondrial complex I in HCT-116 cells ^[59]^. The blockage of complex I is crucial for metformin’s anticancer activity, where cells decrease their proliferation rate under glucose-supplied conditions; however, no change in cell viability is observed. Additionally, depriving cells of glucose triggers a significant increase in cell death ^[59]^. Similarly, our experiments show that HCT-116 exposure to compound **1** decreases cell proliferation at concentrations as low as 0.1aM under typical availability of glucose, while maintaining cell viability (Figures 3A and 3B). According to other studies, the disruption of energy supply caused by blockage of the mitochondrial complex I is sufficient to cease or decrease cell proliferation in the presence of glucose; however, the cells remain active with basal metabolic activities due to energy generation through the glycolytic pathway ^[48,59,60]^. Furthermore, a reduction in the cell proliferation rates, mainly in the HCT-116 cell line, supports the results obtained by the MTT assay despite possible disturbances in MTT-reduction reactions, and, therefore, underestimation of IC_50_ values. Further studies should explore the translational potential of piericidins, particularly considering their application as pharmacological tools and sensitizing agents in cancer therapy. Assessing their efficacy in combination with standard chemotherapeutics, along with their safety and selectivity in preclinical models, will be fundamental to ensure that piericidins remain viable contenders in the path toward drug development.

## Conclusions

*Palythoa variabilis* from Brazilian tropical waters harbors bacteria capable of producing bioactive metabolites. Herein, piericidin A1, glucopiericidin A1 and piericidin C1 were isolated from *Streptomyces* sp. BRA-035, an Actinomycetota strain associated with *P. variabilis*. Piericidin A1 exhibits a highly potent antiproliferative activity at sub-femtomolar concentrations against various cancer cell lines, especially towards cells that favor an oxidative metabolism for energy supply. These findings reinforce the relevance of cnidarian-associated microbiota as a promising source of bioactive natural products.

## Experimental

### Sampling of zoantharians

Colonies of *Palythoa variabilis* were sampled during low tide at Taíba and Paracuru beaches in tide pools of beachrocks (3°30’20.46” S, 38°53’54.31” W and 3°39’5” S, 39°01’3” W respectively), located in the state of Ceará on the tropical northeast coast of Brazil, using surgical forceps and a stainless-steel spatula rinsed with 70% ethanol. Polyps were rinsed with ethanol 70% following wash in sterile sea water. Sampled material was stored in sterile sample bags, packed in styrofoam boxes cooled with icepacks and taken for laboratory processing within 3 hours. Authorization and Information System (SISBIO number 48522-2, SisGen number AC0781C).

### Isolation of bacterial strains associated with zoantharians

The colonies of zoantharians sampled were processed under sterile conditions as follows: samples were cut extensively, heated to 56°C for 15 min, spread on agar-media plates and incubated at room temperature (approximately 25 °C) for up to 8 weeks. From 2 to 12 weeks incubation, 10 strains have grown and were isolated. For cultivation of bacteria associated with *P. variabilis*, the following media were used: SWA (Sea Water Agar - 18 g of agar; 1L of filtered seawater diluted 75% in dH_2_0), TM (Trace Minerals Agar - 0.1 g/L glucose; 0.1 g/L yeast extract; 0.5 g/L K_2_HPO_4_; 0.7 g/L Na_2_HPO_4_; 0.1 g/L KNO_3_; 0.3 g/L NaCl; 0.1 g/L MgSO_4_.7H_2_O; 0.02 g/L, CaCl_2_.2H_2_0; 18 g/L agar; 1L seawater diluted 75% in dH_2_0) and SCA (Starch-Casein Agar - 10 g/L soluble starch; 1 g/L casein powder; 37 g/L seawater preparation; 15 g/L agar; 1 L dH_2_O). Isolation of strains was carried using A1 agar media (10 g/L soluble starch; 4 g/L yeast extract; 2 g/L peptone; 18g/L agar; 1 L filtered seawater diluted 75% in dH_2_0). Purified bacteria strains were grown in liquid A1 broth to produce the crude extract, to deposit in a bacterial bank (MicroMarin) at-80°C in 25% glycerol and for genetic extraction for molecular identification. Isolated strains were cultivated for 8 days (26 ±1°C, under constant agitation) and extracted with AcOEt (Synth) (1:1, during 2 hours under agitation). Solvent was removed by reduced pressure rotary evaporation.

### BRA-035 molecular identification by 16S rRNA sequencing

Genomic DNA from strain BRA-035 was extracted using the DNeasy Blood & Tissue Kit (QIAGEN, Germany) following the manufacturer’s instructions. PCR amplification of the 16S rRNA gene was performed using the Illustra Ready-To-Go RT-PCR Beads kit (GE Healthcare Life Sciences, USA) and the universal primers F27 and R1492. PCR products were visualized by agarose gel electrophoresis, quantified using a NanoDrop spectrophotometer (NanoDrop 2000c/2000 UV-Vis, Thermo Scientific, USA), and purified with the MiniElute PCR Purification Kit (QIAGEN, Germany). Sequencing was carried out by the Sanger method at Macrogen Inc. (Republic of Korea). The chromatograms were trimmed, assembled, and the consensus sequence was generated in Geneious Prime (Biomatters Ltd., New Zealand). Molecular identification was performed through comparative analysis of the BRA-035 16S rRNA gene sequence using the BLAST (Basic Local Alignment Search Tool) database hosted by NCBI. Closely related sequences were retrieved from the EzBioCloud platform (https://www.ezbiocloud.net) and aligned in UGENE (Unipro, Russia); the *Micromonospora carbonacea* sequence (GenBank accession NR_037043) was included as the outgroup for phylogenetic analyses. The multiple sequence alignment file was then used to construct a phylogenetic tree using the RAxML-HPC BlackBox tool available through the CIPRES Science Gateway platform (San Diego Supercomputer Center, USA). The resulting tree was visualized and edited in iTOL version 6.9.0 (Interactive Tree of Life; Letunic and Bork, 2021). The BRA_035 sequence was submitted to the NCBI GenBank platform database, and can be found under the code PX514930.

### Bioassay-guided fractionation and isolation of piericidin A1

The *Streptomyces*-BRA035 culture broth (10L) yields 150 mg of extract (EAcE-035). The fractionation direct injection was performed by high-performance liquid chromatography (HPLC) using a Shimadzu UFLC system equipped with SPD-M20A diode array UV-Vis detector. Separations were performed using a phenomenex® reversed-phase column (250 x 4.6 mm, i.d. 5 μm). The solvents used for the analyses consisted of water (solvent A) and acetonitrile (solvent B) transported at 35°C, with an injection volume (loop) of 200 μl and a flow rate of 4.72 mL/min (5-95% A 0-30 minutes and 100% B 30-40 minutes). The scanning wavelength range of the PDA was set at 190-400nm and chromatograms were recorded between 210-400nm. Results in seven fractions (F1-P8) were tested for cytotoxicity effect in order to guide the isolation of the active substance. The identification of piericidin A1 (PA1) (F8; 1.0 mg) and derivatives present in fraction F7 (0.5 mg), was performed through the interpretation of high-resolution electrospray ionization mass spectra (HRESIMS) acquired using a liquid chromatography-mass spectrometry ion trap and time-of-flight (LCMS-IT-TOF, Shimadzu) spectrometer, consisting of an UFLC (ultra fast liquid chromatography) system coupled to an IT-TOF mass spectrometer equipped with electrospray ionization (ESI) source operating either in positive or negative mode. The mass spectra were recorded in the range of m/z 100-1000 Da, using a potential of 4.0 kV on the capillary, and nitrogen as the desolvation gas. Chromatographic runs were accomplished on a BEH C_18_ column (150 mm × 2.1 mm, 1.7 µm) at 40 °C, injecting 2 µL of samples maintained at 20 °C. The mobile phase is composed of 0.1% formic acid in water (A) and 0.1% formic acid in acetonitrile (B) at a flow rate of 0.4 mL/min (gradient system 5-95% A 0-30 minutes and 100% B 30-40 minutes). As well, nuclear magnetic resonance (NMR) spectrum of hydrogen (^1^H) was obtained either on a Bruker Avance DRX-500 (500 MHz to ^1^H). Chemical shifts are given relative to CDCl_3_ at 7.27 ppm.

### Cell culture and MTT assay

Crude extracts, bioguided-fractionation and **1** were tested against PC-3/M (metastatic prostate carcinoma) cell lineage, while **1** cytotoxicity potential was also evaluated against PC-3 (prostate carcinoma), HL-60 (promyelocytic leukemia), OVCAR-8 (ovarian carcinoma), HCT-116 (colorectal carcinoma), SF-295 (glioblastoma) and B16-F10 (murine melanoma) cell lines. Lineages were acquired by the American Type Culture Collection (ATCC) and grown in RPMI-1640 (Thermo-fisher Sci.) medium supplemented with 10% fetal bovine serum, 2 mM glutamine, 1000 U/mL streptomycin and 100 μg/ml penicillin (Sigma-Aldrich Co.), in a controlled atmosphere of 5% CO2 at 37°C. For MTT assay, cells were seeded into 96-well plates at 5×10^4^ cell/mL 24h prior to addition of samples. Extracts, fractions and **1** were diluted in filtered-sterile Dimethyl Sulfoxide (DMSO - Synth). Control groups received the same amount of DMSO and Doxorubicin (Sigma-Aldrich Co.) (0.01 to 5.0 μg/ml) was used as positive control. With 3 hours remaining to complete the incubation time for each experiment, medium was replaced with fresh medium containing 0.5 μg/ml of MTT (Sigma-Aldrich Co.), and incubated again to fulfill 72 hrs. Media were removed and dried formazan crystals were diluted in 150 μl of DMSO. A plate spectrometer reader (DTX 880 Multimode Detector, Beckman Coulter, Inc. Fullerton, CA, USA) was used to measure absorbance at 570 nm. IC_50_ and CI95 parameters were determined by non-linear regression using GraphPad Prism v9.0. Crude extracts screening concentration was 50 μg/mL. For dose-response investigations BRA-035 crude extract ranged from 0.0031 to 50.0 μg/mL, BRA-035 large scale crude extract ranged from 0.00005 to 10.0 μg/mL, bioguided-fractions ranged from 50 to 0.016 μg/mL, F7 ranged from 1 to 1.02^E-07^ μg/mL, **1** ranged from 1.01^E-15^ 12 μM in PC-3, PC-3/M, OVCAR-8, HCT-116, SF-295, B16-F10 and from 5.74^E-06^ 12 μg/mL in HL-60.

### Cell counting and viability by flow cytometry

HCT-116 and B16-F10 cells were seeded in 24-well plates, treated with **1** at 0.1aM, 14fM, 24pM, 3.8nM and 12μM concentrations and incubated for 24, 48 and 72h. After incubation, cells were trypsinized, resuspended in 500 μL of media and centrifuged at 1500 rpm for 2 minutes (Hettich, model Universal 320R). Pellets were resuspended in 250 μL of a 5 μg/mL PI solution (Sigma Aldrich) diluted in phosphate-buffered saline (PBS). After 15 min of incubation, samples were washed with PBS to remove excess PI and analyzed on a BD Accuri C6 flow cytometer (BD). 10.000 events were counted from each sample, excluding debris and doublets. The cells were arranged according to their linear dispersion, volume, and granularity. Percentages of cells with intact or disrupted plasma membranes were analyzed using markers to delimit cell population at regions with lowest and highest fluorescence, compared with negative control, using the BD Accuri TM C6 Software v.1.0.264.21. Data were analyzed based on the mean and respective standard errors of 3 independent experiments, performed in triplicate on GraphPad Software v9.0. Significant differences between treatment groups were verified by analysis of variance (ANOVA) followed by Dunnett’s test with a significance level of at least 5% (p < 0.05).

## Supplementary Material

Supporting information for this article is available on the **WWW** under http://dx.doi.org/10.1002/**MS-number**.

## Acknowledgements

We thank the funding agencies that made this work possible: Coordenação de Aperfeiçoamento de Pessoal de nível Superior - CAPES and Conselho Nacional de Pesquisa e Desenvolvimento - CNPq.

## Author Contribution Statement

Bianca Del B. Sahm, conceptualization, investigation, data curation, formal analysis, original draft writing, writing-review and editing; Katharine G. D. Florêncio, investigation, data curation, formal analysis, original draft writing, writing-review and editing; Francisco das Chagas L. Pinto, investigation, data curation, formal analysis, original draft writing; Ana I. V. Maia, investigation; Carlos A. M. Rocha, investigation; Paula C. Jimenez, investigation, formal analysis, writing-review and editing, Otília D. L. Pessoa, investigation, data curation; Tito M. C. Lotufo, data curation, writing-review, Leticia V. Costa-Lotufo, conceptualization, project administration, funding acquisition, resources, writing-review and Diego V. Wilke, conceptualization, investigation, data curation, project administration, funding acquisition, resources, original draft writing, writing-review and editing.

## Entry for the Graphical Illustration

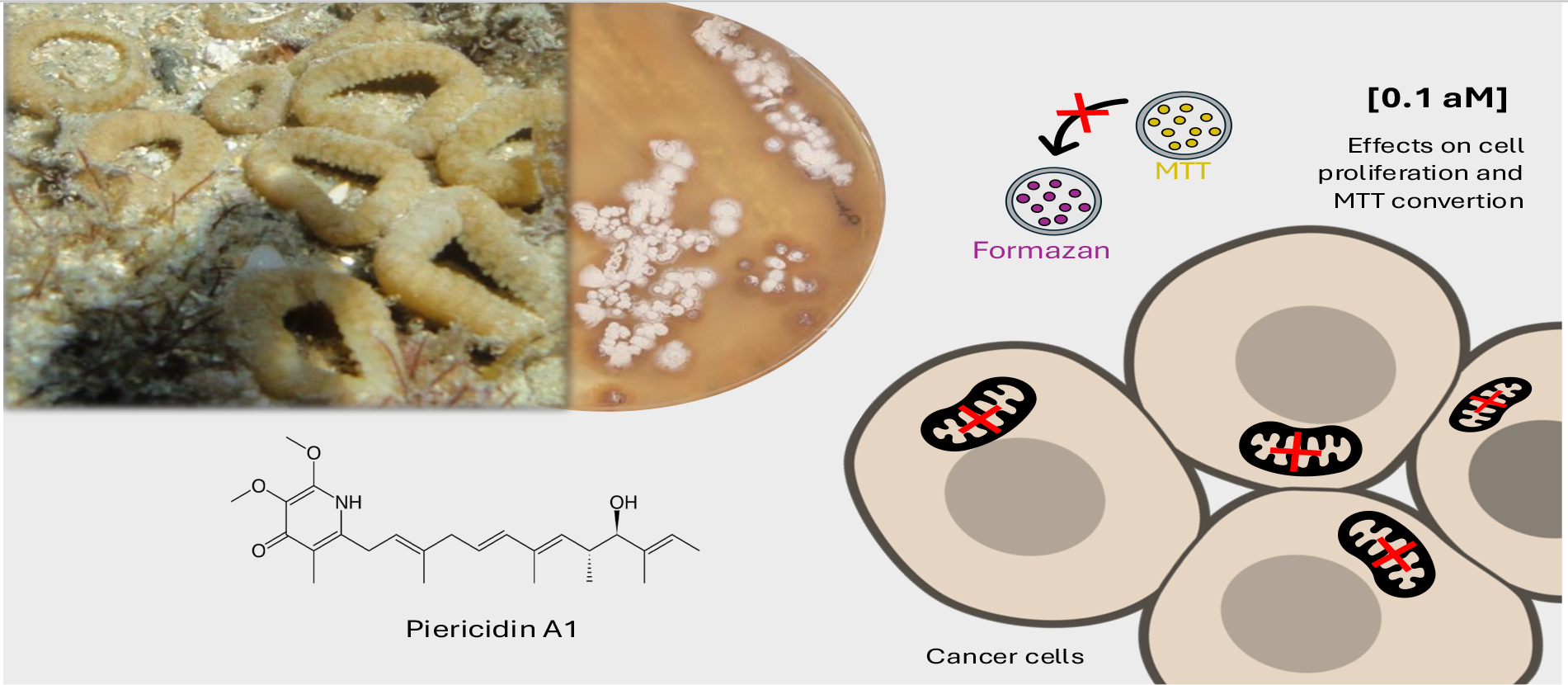

A *Streptomyces* spp. strain, isolated from the zoantharian *Palythoa variabilis*, produces the complex-I mitochondrial inhibitor piericidin A1 with effects on cancer cells proliferation and MTT-formazan reduction at concentrations up to 0.1 aM.

