## Supplementary material for "Anticancer potential of piericidin A1 and derivatives isolated from *Streptomyces* sp. associated with *Palythoa variabilis* from Brazilian Reefs": Suplementary Information

### Supplementary Information

**SI.1.** Bacteria recovered from the association with *Palythoa variabilis* and their respective isolation culture medium.

| Strain | Isolation culture media |
|---|---|
| BRA-035 | SCA |
| BRA-036 | SCA |
| BRA-045 | SCA |
| BRA-046 | SCA |
| BRA-060 | SWA |
| BRA-061 | SCA |

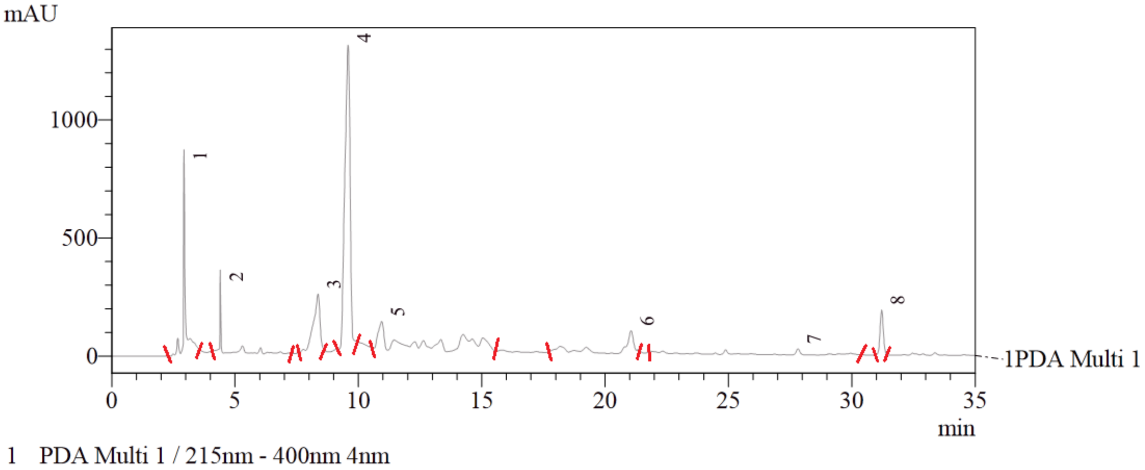

**SI.2.** HPLC chromatogram of the *Streptomyces* spp. (BRA-035) crude extract. Red bars show the delimitation of fractions (numbered) collected through retention time.

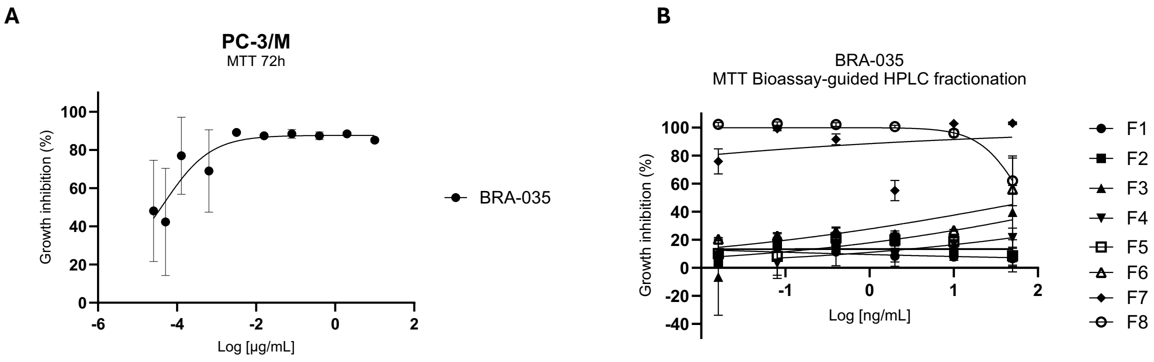

**SI.3.** Bioassay-guided fractionation of BRA-035 crude extract. A- Cytotoxicity profile of BRA-035 large-scale crude extract and B- HPLC-resulted fractions against PC-3/M cell line after 72h of incubation through MTT assay. Analysis was conducted in GraphPad Prism v.10.0 and represent mean±SEM of one experiment performed in triplicate.

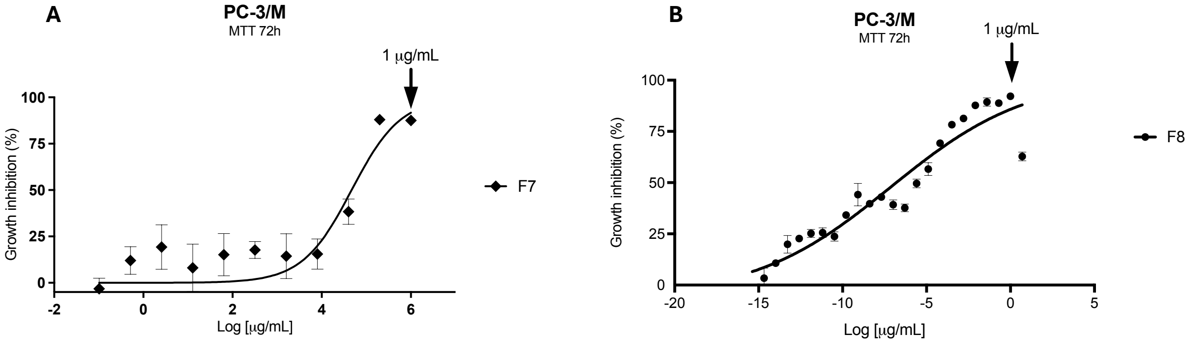

**SI.4.** Cytotoxicity profile of fractions F7 (A) and F8 (B) against PC-3/M cell line after 72h of incubation through MTT assay. Analysis were conducted in GraphPad Prism v.10.0 and represent mean±SEM of one experiment performed in triplicate.

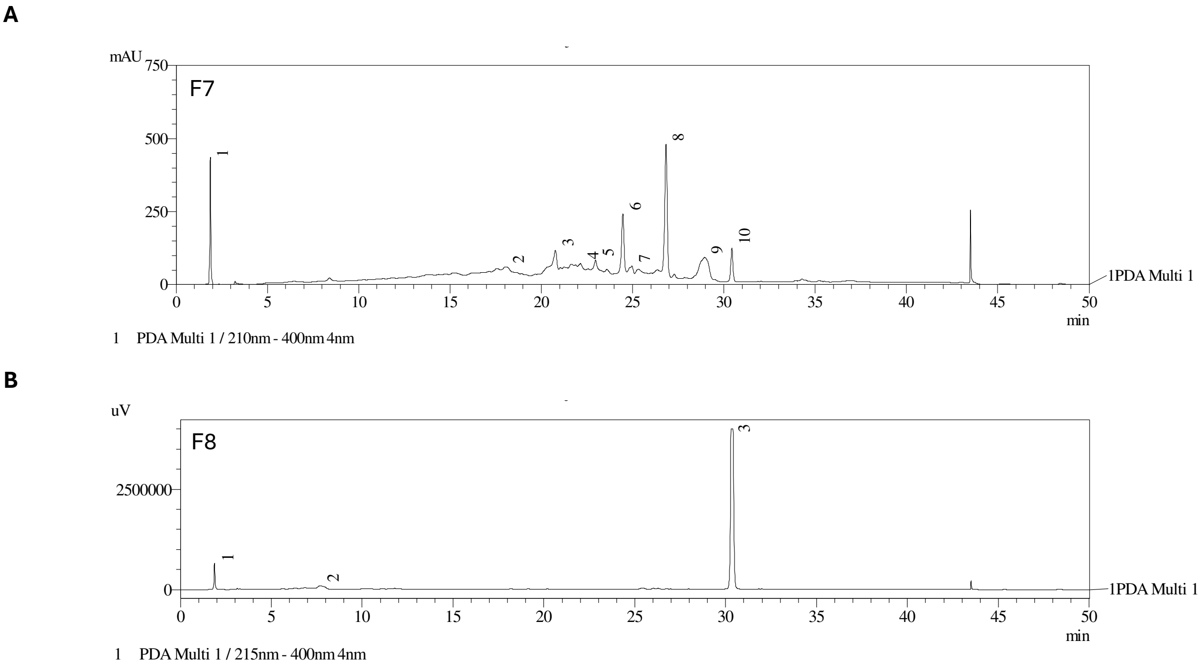

**SI.5.** LCMS-IT-TOF chromatogram PDA Ch1 at 215-400 nm of the fractions BRA035-F7 (**A**) and BRA035-F8 (**B**).

**SI.6.** BRA035-F7 and BRA035-F8 LCMS-IT-TOF chromatogram data.

| Sample | Peak # | Retention time (min) | Area | Height | Area % | Height % |
|---|---|---|---|---|---|---|
| BRA-035-F7[a] | 1 | 1.868 | 1660716 | 435730 | 8.312 | 30.148 |
| | 2 | 18.070 | 5968958 | 33018 | 29.876 | 2.284 |
| | 3 | 20.768 | 1270632 | 67754 | 6.360 | 4.688 |
| | 4 | 22.967 | 567464 | 19377 | 2.840 | 1.341 |
| | 5 | 24.960 | 542973 | 33756 | 2.718 | 2.336 |

| | | | | | | |
|---|---|---|---|---|---|---|
| | 6 | 26.828 | 1868159 | 201431 | 9.351 | 13.937 |
| | 7 | 28.940 | 346169 | 19486 | 1.733 | 1.348 |
| | 8 | 26.828 | 4251490 | 445478 | 21.280 | 30.822 |
| | 9 | 28.940 | 2553391 | 73483 | 12.781 | 5.084 |
| | 10 | 30.434 | 948823 | 115800 | 4.749 | 8.012 |
| Total BRA-035-F7 | - | - | 19978775 | 1445313 | 100.000 | 100.000 |
| BRA-035-F8[b] | 1 | 1.872 | 2576641 | 646406 | 4.950 | 13.795 |
| | 2 | 7.682 | 1723818 | 64872 | 3.312 | 1.384 |
| | 3 | 30.304 | 47748492 | 3974983 | 91.738 | 84.822 |
| Total BRA-035-F8 | - | - | 52048951 | 4686261 | 100.000 | 100.000 |

[a] PDA Ch1 210nm - 400nm 4nm . [b] PDA Ch1 215nm - 400nm 4nm.

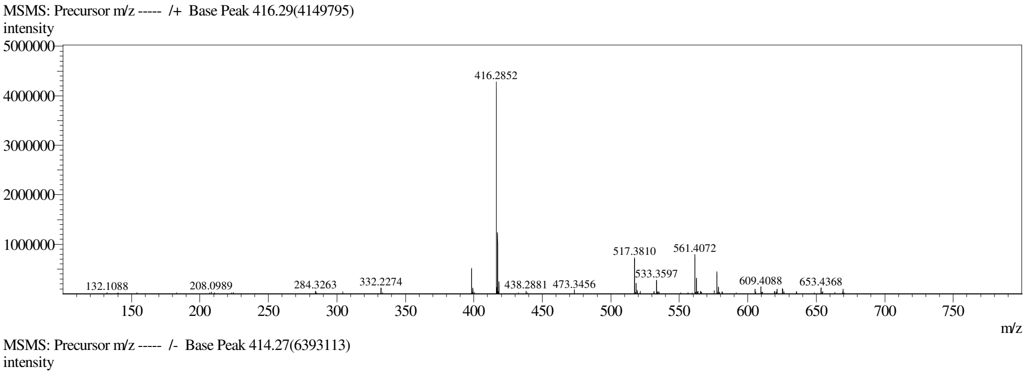

**SI.7.** Fraction F8 LCMS-IT-TOF spectrum for pieridicin A1 (**1**)

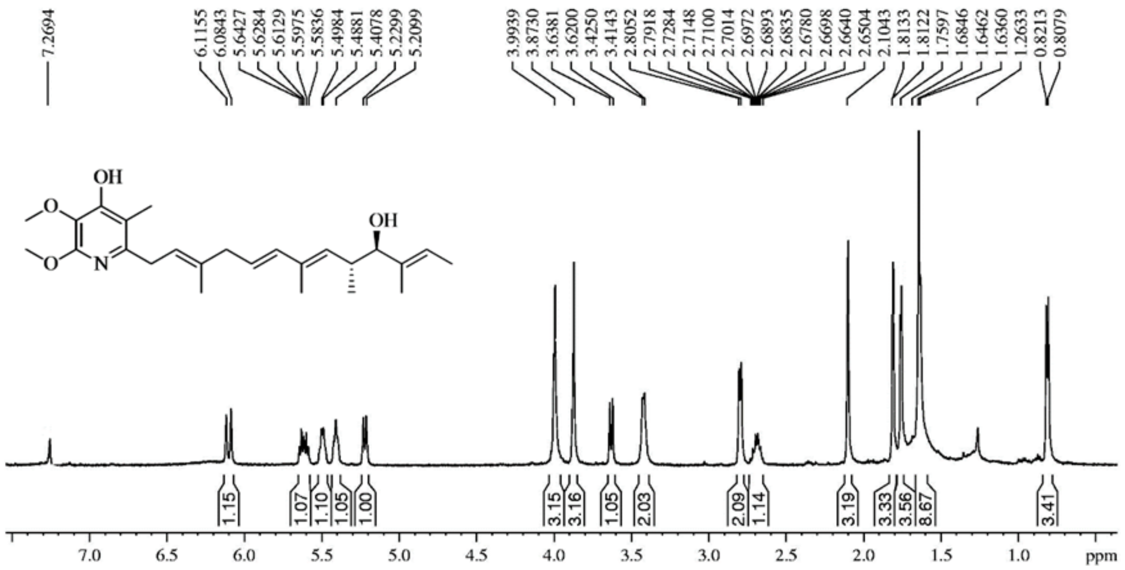

**SI.8.** $^{1}$H NMR in CDCl3 spectrum for pieridicin A1 (**1**)

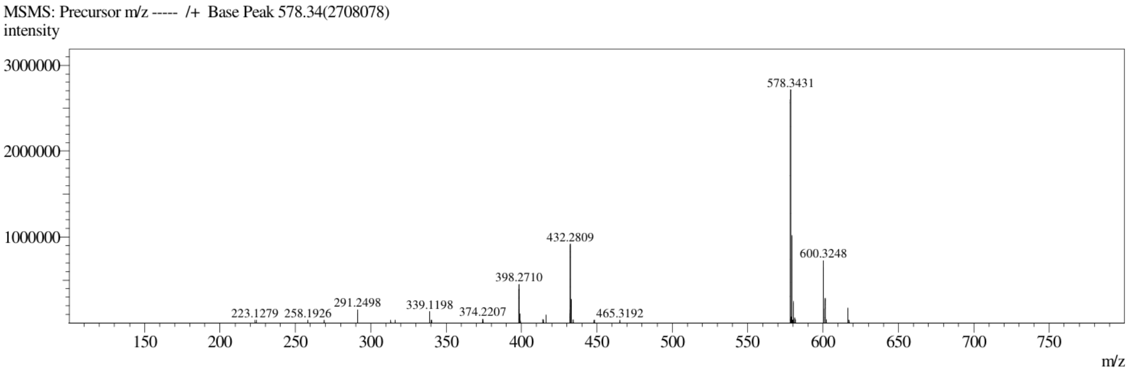

**SI.9.** Fraction F7 LCMS-IT-TOF spectrum for glucopiericidin A1 (**2**).

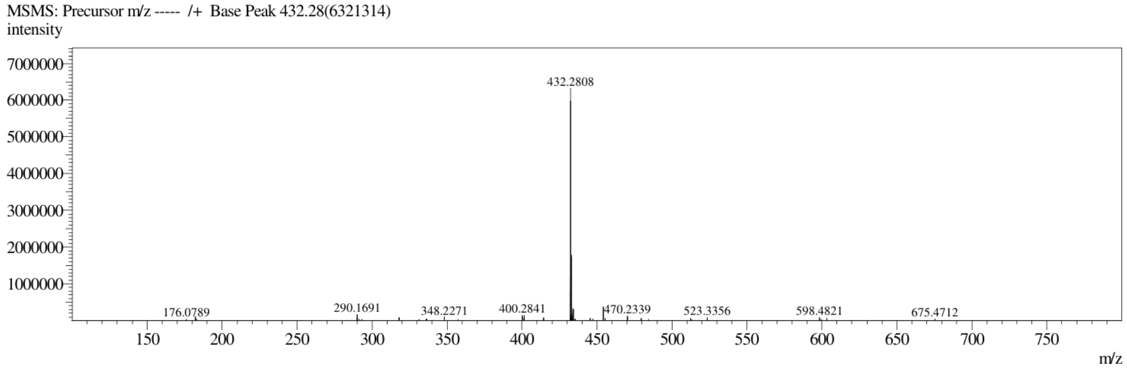

**SI.10.** Fraction F7 LCMS-IT-TOF spectrum for piericidin C1 (**3**).

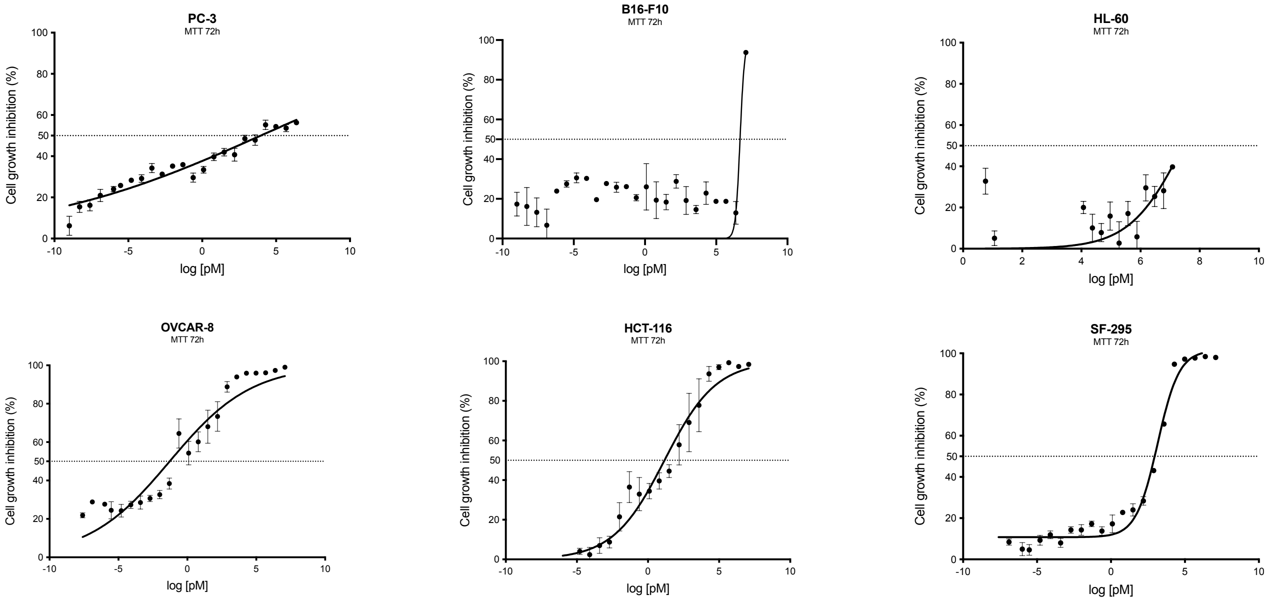

**SI.11.** Concentration-effect graphs of tumor cells incubated with (**1**) for 72h by the MTT assay. Analysis was conducted in GraphPad Prism v.10.0 and represent mean±SEM of one or three experiments performed in duplicate or triplicate.
